# Historic Genomes Uncover Demographic Shifts and Kinship Structures in Post-Roman Central Europe

**DOI:** 10.1101/2025.03.01.640862

**Authors:** Jens Blöcher, Leonardo Vallini, Maren Velte, Raphael Eckel, Léa Guyon, Laura Winkelbach, Mark G. Thomas, Nadia Gharehbaghi, Cassandra T. Mitchell, Jonas Schümann, Sophie Köhler, Elsa Seyr, Katharina Krichel, Sophie Rau, Jana Hirsch, Jana Duras, Kristin Klement, Miriam Wilkenhöner, Lisa Vetterdietz, Francesca Gentilin, Melany Müller, Anna-Lena Mücke, Nicoletta Zedda, Youssef Tawfik, Eveline Saal, George McGlynn, Barbara Bramanti, Jörg Orschiedt, Regina Molitor, Barbara Fliß, Ines Spazier, David Shankland, Claus Vetterling, Kurt Karpf, Vera Planert, Stefan Hölzl, Silvia Codreanu-Windauer, Dieter Quast, Ilija Mikić, Bernd Päffgen, Maxime Brami, Thomas Richter, Raphaëlle Chaix, Susanne Brather-Walter, Peter Steffens, Markus Marquart, Thomas Becker, Jochen Haberstroh, Mischa Meier, Sebastian Schmidt-Hofner, Sebastian Brather, Michaela Harbeck, Steffen Patzold, Daniel Wegmann, Joachim Burger

## Abstract

Many European towns and villages trace their origins to Early Medieval foundations. In former Roman territories, their emergence has traditionally been linked to mass migrations from outside the Roman Empire. However, recent studies have emphasised local continuity with some individual-level mobility. We generated and analysed 248 historic genomes from Late Roman (3rd and 4th century CE) and Early Medieval (5th–8th century CE) burial sites in southern Germany, comparing them to over 2,500 contemporary and Iron Age genomes in addition to 1,344 modern-day genomes from Germany, Italy and Great-Britain. Despite small inferred Early Medieval period community sizes, genetic diversity exceeded that of modern German cities. In the Altheim graveyard, established in the 5th century by a group of Northern European descent, we inferred a demographic shift in the 6th century with the integration of newcomers with ancestry typical of a nearby Roman military camp, likely as a result of the collapse of Roman state structures. We reconstructed multigenerational pedigrees and, using a novel approach to infer ancestry of unsampled relatives, inferred immediate intermarriage between incoming and local groups, with a distinct tendency for men from former Roman background marrying women of northern descent. Burial proximity correlates strongly with kinship, in some cases spanning six generations. These communities were organized around small family units, exhibited loosely patrilineal or bilateral descent patterns, practiced reproductive monogamy, and avoided close-kin marriages. Such practices reflect broader transformations in family structures that began during the Late Roman period, were transferred to small agrarian societies in the Early Medieval period, and continued to shape European societies. By the 7th century, ongoing admixture had shaped genetic diversity patterns into those resembling Central Europe today.

## Introduction

During the transition from the Late Roman period to the Early Middle Ages (4th to 6th century CE), Central Europe underwent significant political, cultural and demographic changes influenced by the dissolution of Roman rule, the spread of Christianity and the development of new settlement patterns. The political landscape shifted dramatically, with the emergence of new polities, leaving a lasting impact, particularly in the southern part of Central Europe. However, our knowledge about these localities and their societies remains scant. Written records from this period are extremely scarce, and only few Early Medieval settlements have been excavated. Consequently, cemeteries associated with these settlements have become invaluable historical sources. Distinctive furnished burials, traditionally described as “row graves”, began to appear in the former Roman imperial border regions across Europe from the mid-5th century onwards ^1^. In addition to skeletal remains, these graves frequently contain artefacts such as clothing, weapons, jewellery and vessels, which provide insights into the lives and deaths of ordinary people in Early Medieval Central Europe ^2^. Archaeological and historical research has identified the societies behind the row grave horizon in present-day southern Germany as rural communities of up to a dozen farmsteads that primarily relied on cultivating crops and raising of pigs and cattle ^3^. Nevertheless, they maintained far-reaching contacts and developed social hierarchies ^4,5^. Some graves display Christian symbols as early as in the late 6th century CE ^6^.

To broaden our understanding of the population histories and demographic processes in this shifting landscape, we sequenced 248 ancient genomes (median depth 2.25 X) from multiple archaeological sites located in the northern frontier zone of the Roman Empire (Fig. 1). We focused on Early Medieval row graves from two regions: the Danube-Isar area in Upper and Lower Bavaria, represented by sites like Altheim and Weilheim, and the Rhine-Main area, represented by Büttelborn and Mömlingen (Fig. 1). During Antiquity, both regions were part of the Roman Empire. While the Rhine-Main area belonged to the province of *Germania Superior* that persisted until the end of the 3rd century, the Danube-Isar region belonged to the province of *Raetia Secunda* until the Western Roman Empire collapsed in the late 5^th^ century. It then became part of the Ostrogothic kingdom, though the extent of its control and northern reach remains debated ^7^. By around 540 CE, the Danube-Isar region fell under Frankish influence, with the Franks establishing a military command led by a duke (*dux*), which later evolved into the foundation of the Duchy of Bavaria.

**Fig. 1.**
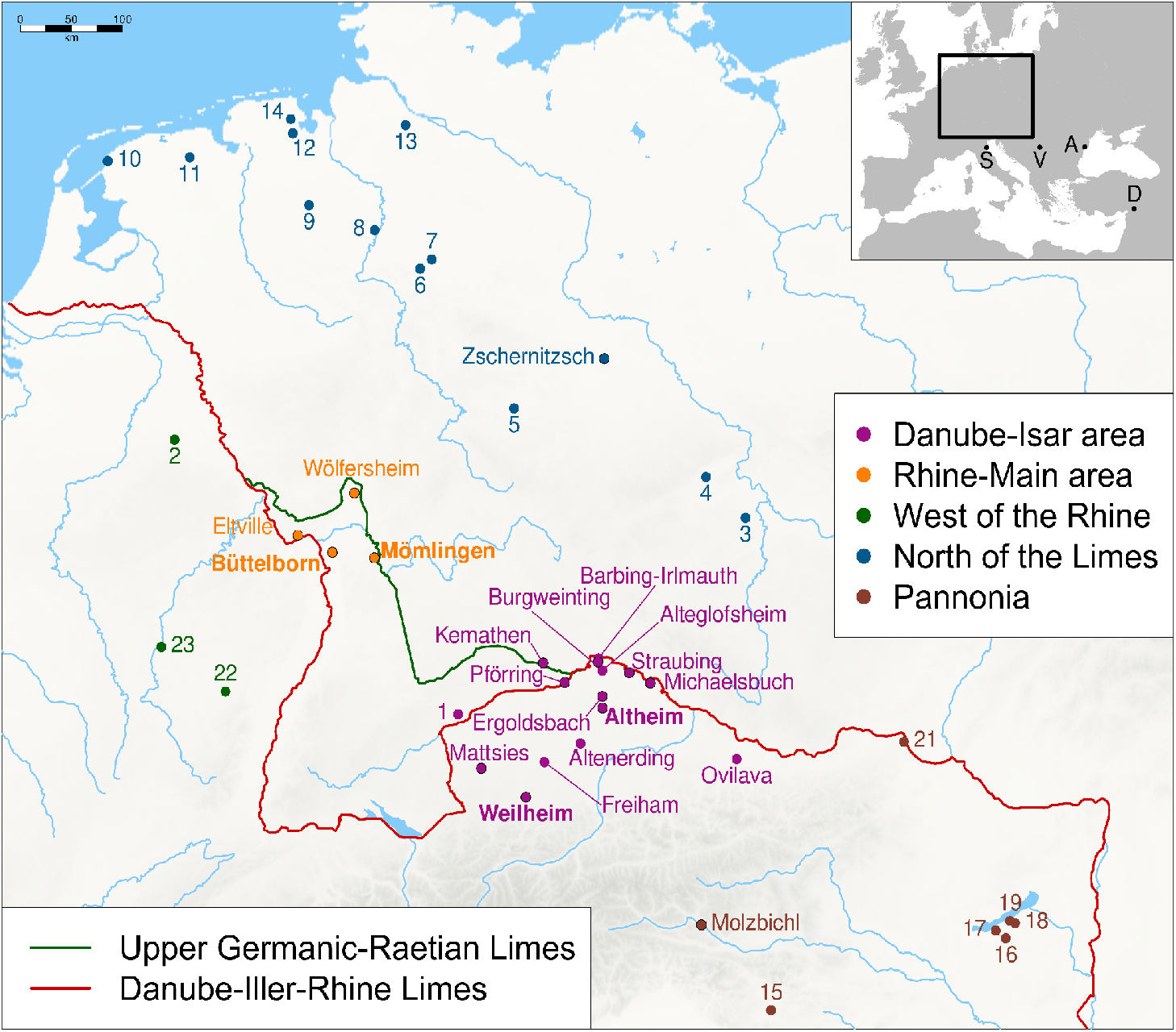
Location of the burial sites examined in this study. The Upper Germanic-Rhaetian Limes, depicted in green, marked the boundary of the Roman Empire until the second half of the 3rd century CE. The Danube-Iller-Rhine Limes, shown in red, delineated the Roman military frontier from the late 3rd century until the early 5th century. Late Roman (Straubing-Azlburg, Pförring, Kemathen) and Early Medieval sites with new data reported in this study display a black border (with the four main ones in bold), while sites from outside the core region are marked by letters: Viminacium (V), Argamum (A), Spina (S), Doliche (D). Published reference sites are numbered: Niederstotzingen (1), Alt-Inden (2), Brandysek (3), Konobrze (4), Hassleben (5), Hiddestorf (6), Anderten (7), Liebenau (8), Drantum (9), Midlum (10), Groningen (11), Zetel (12), Issendorf (13), Schortens (14), Ljubljana (15), Hács (16), Fonyód (17), Szólád (18), Balatonszemes (19), Komárno (20), Klosterneuburg (21), Sarrebourg (22), Metz (20).

For comparative purposes, we supplemented this dataset with new and published Early Medieval genomes from present-day southern and eastern Germany, as well as Austria and Hungary ^8–15^. To explore genomic variability in earlier periods, we generated additional 36 genomes from nearby Late Roman sites, Straubing-Azlburg, Kemathen, and Pförring, as well as key sites across Europe and beyond: Viminacium (Serbia), Argamum (Danube Delta), Doliche (Anatolia), and Spina (Italy) (Fig. 1, SI Chapter S1, External_Data_Table_1).

### Shifting Ancestries, Population Structure and high genetic diversity

We first performed Principal Components Analysis (PCA) to compare individuals from Altheim in the Danube-Isar region — our primary study site — with a dataset of 765 Iron Age genomes from western Eurasia, grouped into clusters based on their geographic origin (External_Data_Table_2.1-2, Fig. S6.4). This analysis revealed three distinct ancestry phases in the Altheim graveyard, indicative of major demographic shifts (Fig. 2A).

**Fig. 2:**
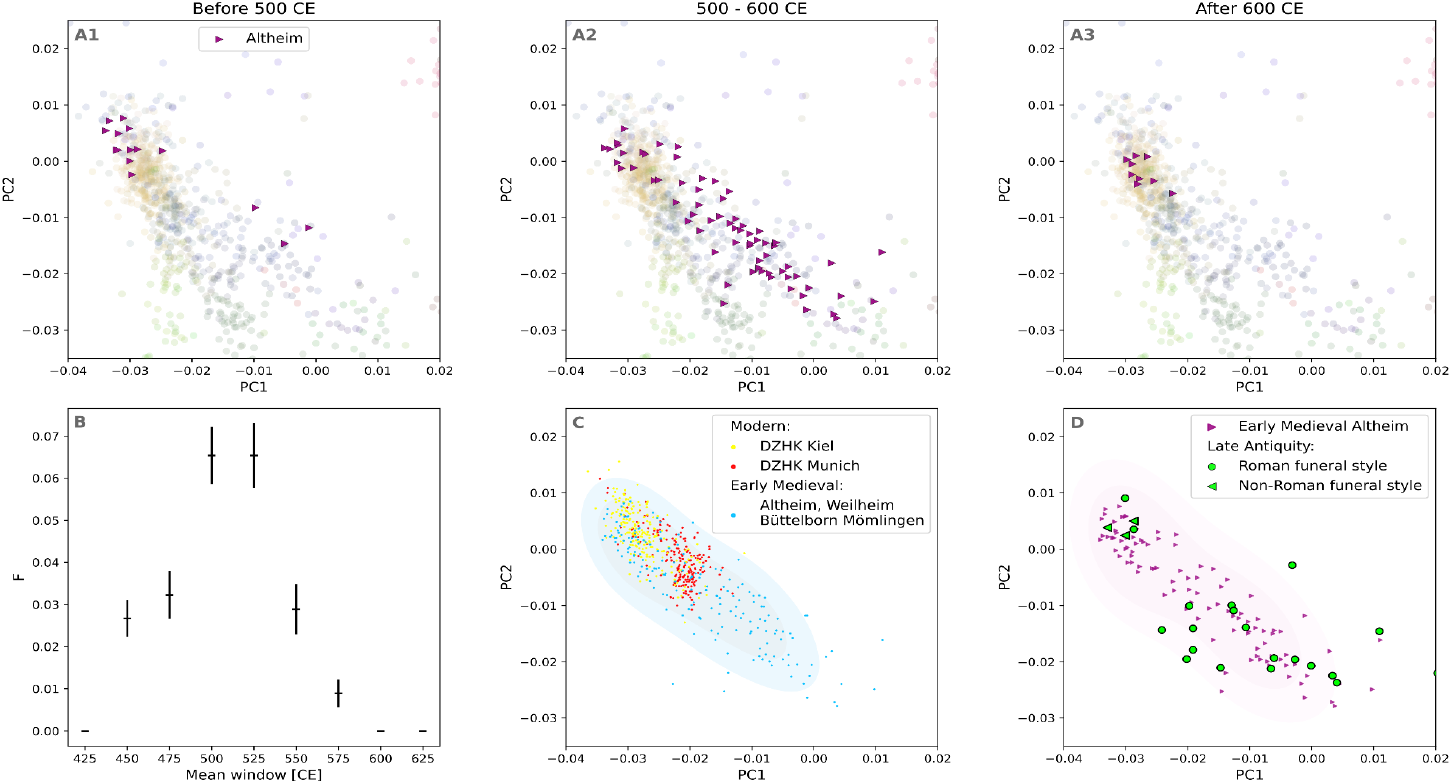
(A1-3): Principal Component Analysis (PCA) of genetic variation in Altheim over time. The chronology is based on estimated individual birth dates. The background reference genomes represent genetic variation across Europe from 800–1 BC. The spatial distribution of these reference samples and the corresponding colour code are provided in Supplementary Information (SI) Chapter S6. **(B): Inbreeding coefficient (F) for the Altheim population**. Estimates of F with standard errors over time, illustrating the development of population substructure across different periods. **(C): PCA of Early Medieval genomes compared to modern individuals sampled at two German hospitals**. Munich (southern Germany) and Kiel (northern Germany), with 75% contour lines. Data were provided by the DZHK (German Centre for Cardiovascular Research). For additional details, see Fig. S6.5-S6.8. **(D): PCA comparing Early Medieval genetic variation in Altheim with genomes from the Late Antiquity of the Danube-Isar region and Upper Austria**. Roman funeral style: Straubing-Azlburg and Ovilava sites, non-Roman funeral style: Pförring and Kemathen sites.

In the earliest phase, individuals born before 500 CE predominantly cluster with Iron Age samples from Northern and Central Europe, consistent with descent from ancestral populations further north. However, individuals born in the 6th century CE exhibit a much broader distribution in PCA space. While some still overlap with Iron Age populations from Northern and Central Europe, others show affinities to groups from the western Mediterranean, Southeast Europe, in six cases the Steppe region, or even fall completely outside the genomic variation of western Eurasia (Fig. S6.2-S6.3). This expanded genetic range indicates a substantial influx of people with diverse ancestries. Individuals from the later phase of occupation, after 600 CE, once again cluster with Northern and Central European Iron Age genomes. However, their position in PCA space shifted slightly toward Southern Europe, suggesting a degree of genetic admixture, likely occurring in the preceding 6th century.

We next assessed population substructure over the duration of these demographic shifts. Population substructure results in a deviation from random mating, which we quantified over time with the inbreeding coefficient (F). During the 5th century, we found low F-values, indicating little population substructure. Shortly after 500 CE, F-values peaked before gradually declining toward the 7th century (Fig. 2B, Table 1). This pattern aligns with the arrival of genetically diverse groups around 500 CE, prior to their assimilation.

**Table 1:** Population genetic statistics of the four main Early Medieval study sites and modern 1000 genomes populations from Great Britain (GBR) and Central Italy (TSI).

|  | Altheim | Büttelborn | Mömlingen | Weilheim | GBR | TSI |
| --- | --- | --- | --- | --- | --- | --- |
| number of individuals | 102 | 40 | 28 | 13 | 88 | 107 |
| % of individuals with a 1st-3rd degree relative | 66.6% - 83.4% | 55.8%-84.2% | 5.9% - 36.1% | 5.9%-56.1% | / | / |
| Est. community size per generation and settlement (5 village model) | 36-64 | 22-53 | / | / | / | / |
| Mean of pairwise shared IBD fragments >8 centimorgan - both sexes | 103.51<br>± 5.59 | 137.49<br>± 18.17 | 24.99<br>± 8.84 | 85.94<br>± 47.29 | 2.36<br>± 24.61 | 1.68<br>± 8.61 |
| % pairs with 0 IBDs >8cM | 77.5 % | 87.58 % | 92.59 % | 91.03 % | / | / |
| Mean pairwise shared IBD fragments >8 centimorgan - all females | 40.03 ± 7.22 | 125.60 ± 27.05 | 10.84 ± 7.47 | 129.74 ± 100.57 | 2.72 ± 28.60 | 1.34 ± 5.40 |
| Mean pairwise shared IBD fragments >8 centimorgan - all males | 170.50 ± 13.19 | 177.74 ± 53.35 | 38.28 ± 19.10 | 0.00 ± 0.00 | 1.47 ± 8.19 | 1.95 ± 7.07 |
| mt haplotype diversity (CI) (unrelated) | 0.99 (0.93 - 1.04) | 0.99 (0.89 - 1.08) | 0.98 (0.91 - 1.04) | 0.97 (0.79 - 1.15) | 0.92 (0.90 - 0.95) | 0.92 (0.88 - 0.96) |
| Y haplotype diversity (CI) (unrelated) | 0.86 (0.51 - 1.20) | 1.00 (0.55 - 1.45) | 0.65 (0.34 - 0.96) | 0.50 (-0.15 - 1.15) | 0.39 (0.16 - 0.63) | 0.75 (0.63 - 0.87) |
| average pairwise difference ($\pi$) (transversions, without closely related) | 5th: 0.0592 ± 0.0007 | 0.0576 ± 0.0007 | 0.0602 ± 0.0007 | 0.0606 ± 0.0007 | 0.0505 ± 0.0007 | 0.0509 ± 0.0007 |
|  | 6th: 0.0592 ± 0.0007 |  |  |  |  |  |
|  | 7th: / |  |  |  |  |  |
|  | total: 0.0597 ± 0.0006 |  |  |  |  |  |
| Genome wide $\theta$ (unrelated) | 0.000853 ± 0.000125 | 0.000925 ± 0.000101 | 0.000803 ± 0.000059 | 0.000866 ± 0.000035 | / | / |
| inbreeding coefficient (F) (without closely related) | 5th: 0.0393 ± 0.0061 | 0.0409 ± 0.0069 | 0.1270 ± 0.0105 | 0.0261 ± 0.0052 | 0.0117 ± 0.0035 | 0.0107 ± 0.0036 |
|  | 6th: 0.0519 ± 0.0071 |  |  |  |  |  |
|  | 7th: / |  |  |  |  |  |
|  | total: 0.1422 ± 0.0104 |  |  |  |  |  |
| F<sub>st</sub> between (transversions) | Altheim - Büttelborn | Altheim - Mömlingen | Büttelborn - Mömlingen | Weilheim - Altheim | Weilheim-Büttelborn | Weilheim-Mömlingen |
|  | 0.0122 ± 3.47E-18 | 0.0080 ± 1.73E-18 | 0.0093 ± 1.73E-18 | 0.0118 ± 1.73E-18 | 0.0141 ± 5.20E-18 | 0.0107 ± 3.47E-18 |
|  | TSI - GBR | TSI - CEU | GBR - CEU | Altheim 5th - 6th century |  |  |
|  | 0.0044 ± 8.67E-19 | 0.0041 ± 8.67E-19 | 0.0017 ± 2.17E-19 | 0.0140 ± 1.73E-18 |  |  |
| inbreeding coefficient (F) over all sites (without closely related) | Early Medieval sites southern Germany |  |  |  | Modern GBR + TSI |  |
|  | 0.1287 ± 0.0051 |  |  |  | 0.0116 ± 0.0029 |  |

We then used qpAdm to model the ancestries of Altheim individuals, using various Iron Age populations as potential sources (SI Chapter S7, External_Data_Table_2.3). Consistent with the patterns identified using PCA and F-statistics, the majority of 5th-century individuals (81%) can be modeled as deriving their ancestry exclusively from Northern and Central European Iron Age populations (hereafter IA-NC). During the 6th century, 51% of genomes are still best described by 100% IA-NC ancestry, while the remaining individuals exhibit partial or complete ancestry from Iron Age populations of the western Mediterranean, the Balkans, or western Anatolia. Since qpAdm modeling revealed no clear genetic differentiation among Iron Age groups from the more southern sources — often identifying them interchangeably as potential sources of ancestry — we collectively refer to them as “Iron Age - Southeast” (hereafter IA-SE). This designation is based on the observation that Altheim individuals, in PCA space, display a stronger genetic affinity to Iron Age and Late Antiquity genomes from south-eastern Europe, such as those from Argamum, than to those from Italy and the western Mediterranean, such as the Spina site (see Fig. S1.19, S1.21). Additionally, five individuals show minor genetic contributions from Steppe-related populations, and one individual (Alh_245) has approximately 75% East-Asian-like ancestry (Iron Age populations from China Xianbei; Mongolia Khentii-Xiongnu, External_Data_Table_2.4-5). By the final 7th century, 94% of the individuals can be modeled as deriving all their ancestry from IA-NC — consistent with further genetic contributions from northern populations.

Nearly all Early Medieval cemetery population samples in southern Germany exhibit high diversity across all measurable indices, including π, θ, mitochondrial and Y-chromosomal haplotype diversity, and PCA (Table 1, Table S5.1, Fig. S6.2-S6.3). The average pairwise differences (π) estimated for Altheim, for instance, exceed that of the 1000 genomes present-day reference populations (Table 1). This aligns with the observation that Early Medieval individuals from Altheim (as well as Büttelborn, Mömlingen and Weilheim) are spread across a significantly larger PCA space than individuals of modern German cities (Munich and Kiel^16^, Fig. 2C, Fig. S6.5-6.8). Notably, the diversity (e.g. π) observed in Altheim was consistently high in both the 5th and 6th centuries, suggesting that the incoming groups contributed additional genetic diversity to an already genetically diverse *in-situ* population.

### Similar demographic processes across southern Germany

While the smaller sample sizes at other archaeological sites do not allow the same fine-grained chronological analysis as in Altheim, we do find evidence of similar processes occurring throughout the region. Individual affinities towards IA-NC-like and IA-SE-like ancestries quantified with D-statistics (Fig. S8.1, External_Data_Table_2.6), for instance, confirmed an overall influx of IA-SE-like ancestry across the region, although there was variation in both timing and magnitude across sites, apparently influenced by the proximity to functioning Roman settlements (Fig. S8.2, Supplementary Video S1). In the Rhine-Main region, for instance, the five genomes of Wölfersheim represent the earliest period (around 500 CE) and all fall within the IA-NC range, with one individual showing some additional Iron Age Steppe-like ancestry. The available genomes from Eltville, located outside the former Roman frontier but close to the large Roman settlement of *Moguntiacum* (today Mainz), are from the 6th century and represent the full spectrum from IA-NC to IA-SE. Interestingly, that signal is much weaker and appears 2-3 generations later at Büttelborn and Mömlingen located further away from the former Roman frontier (Fig. S8.1).

The studied Early Medieval settlements from the Danube-Isar area consistently exhibit the full spectrum from IA-NC to IA-SE ancestry, whenever sufficient data for the 6th century is available, reflecting their location within a region that remained under Roman control until the end of the 5th century. At the southernmost site of Weilheim, furthest from the Limes (a Latin term for ‘border’), individuals with appreciable IA-SE ancestry are observed as early as 500 CE. A comparable diversity is evident in contemporaneous genomes from Straubing-Bajuwarenstrasse ^8^, a cemetery adjacent to the Late Roman burial site of Straubing-Azlburg (3rd and 4th century), itself associated with a Roman fortress. In Burgweinting, a site directly at the Danube-Limes 50 km north of Altheim and linked to it through several longer identity by descent (IBDs) segments (Fig. 3, Fig. S9.4-S9.7, External_Data_Table_3), includes an elite group burial showing similar signs of a demographic shift during the 6th century. Unlike in Altheim, however, individuals with different ancestry backgrounds were buried separately (Fig. S1.9). Taken together, increased diversity during the 6th century appears to have been particularly pronounced in the hinterlands of former Roman territories and near Roman settlements, whereas it developed more gradually in more peripheral regions.

**Fig. 3:**
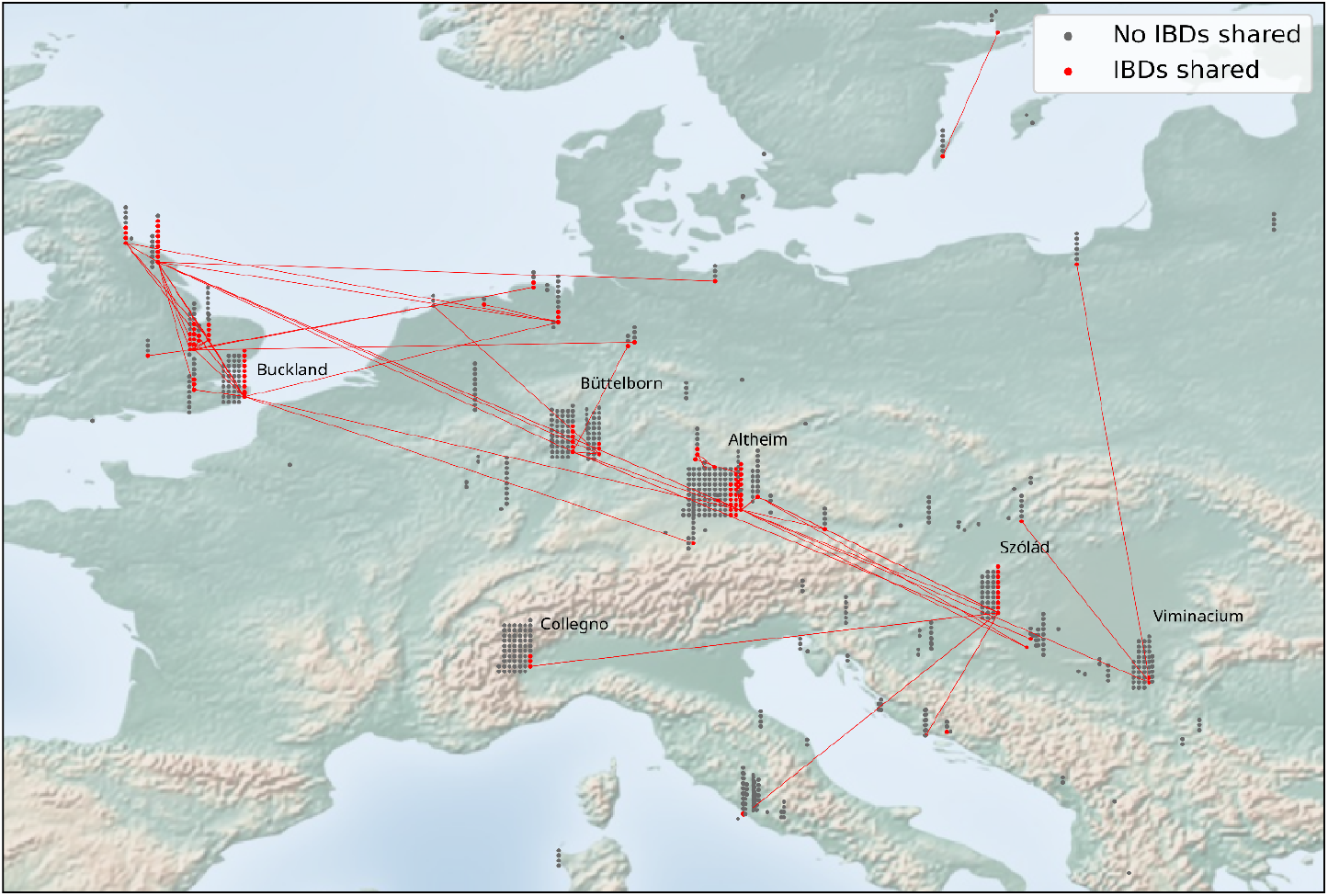
Map showing pairs of individuals from Late Antiquity and the Early Middle Ages who share long genomic segments identical by descent (IBD). Sites where individuals are sharing at least one segment of 20 centimorgans are linked by a red line. The vast majority of IBDs are shared between individuals with an IA-NC-like ancestry profile (see also Fig. S6.13). The sites with the highest number of individuals in each area are indicated by name.

### Northern Ancestry Connects European Regions

Remarkably, 14 Early Medieval individuals from the Rhine-Main and Danube-Isar regions share long IBD segments (>20 cM) with contemporaneous individuals from distant regions such as northern Germany and England, as well as Austria, Hungary, Croatia, and Serbia — including the Roman city of Viminacium (Fig. 3, External_Data_Table_3). In our dataset, long IBD segment sharing is almost exclusively present among individuals who appear in the upper (i.e. IA-NC-like) region of the PCA plot (Fig. S6.13). The frequency of such long-distance connections generally decreases both over time and in association with individuals whose genomes appear toward the lower region of the PCA plot (Fig. S9.8-S9.14). These patterns suggest that the northern groups in our Early Medieval sites were more closely interconnected, likely tracing their ancestry to a network of groups who resided predominantly well north of the Roman frontier during preceding centuries.

We have identified a pair of likely second cousins buried more than 270 km apart in Büttelborn (Rhein-Main area) and Hiddestorf, a small burial ground associated with local elites in Lower Saxony ^10^. Furthermore, there are significant biological kinship connections within the Danube-Isar region that extend to an esteemed kin group in the Pannonian site of Szólád, situated approximately 500 km away ^9^. Two young siblings from Szólád (SZ8, SZ14) are third or fourth degree related to a male individual from Altenerding (AED92b) through their mother, while their paternal lineage is connected to two siblings in Straubing-Bajuwarenstrasse (STR355c, STR491; 5th or more distant degree ^8^). Additionally, an individual from Szólád (SZ13) is connected by fifth or sixth degree to a female from Altheim (Alh_158), who herself shares third- or fourth-degree kinship with AED92b. These findings suggest that a portion of the northern population in the Early Medieval Danube-Isar sites had close ties to the group that arrived in Pannonia around the middle of the sixth century (SI Chapter S9). Most of these identified relationships do not indicate active biological kinship or marriage networks during the lifetimes of the Early Medieval communities, as the shared ancestors likely predate the period under study. However, the ancestral ties between Szólád and the Danube-Isar region endured until shortly before the establishment of the cemeteries in southern Germany, so that the common ancestors were surely still remembered.

### Roman Borderlands: Hub of Attraction

In the following, we define “migration” as the unidirectional relocation of individuals across cultural or political boundaries, while “mobility” refers to omnidirectional, often shorter-distance movements within a shared cultural or political region ^17^. As shown above, 5th-century individuals from Altheim represent only a subset of the genomic diversity seen in southern Germany and neighbouring regions during the Iron Age ^11,18^. Most display strong genetic similarities with Iron Age populations from Northern Europe, such as those from Denmark and the Lowlands, suggesting that by the mid-5th century — or earlier — migrants from the north had begun organizing burials in new cemeteries across the former Limes area. Prior studies support a southward migration from Northern Europe between 1–500 CE, reaching Pannonia by the 5th century ^9,13,19,20^.

Several lines of evidence suggest that much of the migration from northern regions to the provincial borderlands, or even into the Roman Empire, took place before the Early Medieval period. First, 17% of Altheim individuals, all from the IA-NC-like group, share long IBD segments (>20 cM) with individuals of nearby sites (80 km radius, Fig S9.7, Extended_Data_Table3) such as Burgweinting (seven pairs, 26 cM - 62 cM), Straubing-Bajuwarenstrasse (two pairs, 22 cM - 25 cM) and Ergoldsbach (eleven pairs, 21 cM - 498 cM). This includes an Altheim female (Alh_161) who is kin (1st to 2nd cousin) to three brothers from nearby Ergoldsbach, and another female from Burgweinting who is a sister of a woman from Altenerding ^8^. In contrast, no such connections were found between the Danube-Isar and Rhine-Main regions, suggesting genetic isolation for at least eight generations. Second, the rate of shared IBD segments (>8 cM) decreased with geographic distance (r:0.44, p<0.019), consistent with isolation-by-distance rather than recent migration from a homogenous source population (Fig. S9.10). Third, and in line with this interpretation, the combined row-grave population of southern Germany is genetically highly substructured (F=0.129) and shows significantly higher F_ST_ values between settlements than modern continent-wide populations (Table 1). All of this suggests that the mid-5th century populations along the southern German Limes were already genetically differentiated when the cemeteries were established, likely because their ancestors moved to the region some generations or even centuries earlier. The presence of individuals with IA-NC ancestry in the region already during the preceding Late Roman period is directly confirmed both genetically and archaeologically by two individuals from Straubing-Azlburg and three individuals from the chamber graves of Pförring and Kemathen. These burials belong to a very small group of elite special burials, placed in wooden chambers, which occur in the Central European *Barbaricum* during the second half of the 4th and the first half of the 5th century, occasionally extending into southern Germany (Fig. 1, Fig. 2D).

Stable isotope analysis, however, indicates that a significant proportion of individuals from the 5th-century Danube-Isar-area changed their geological environment between childhood and adulthood. Strontium isotope data from over 100 individuals from various Early Medieval sites, dating back to the 5th century, revealed a notably high ‘minimum frequency of migrants’ (MFM) of 23% ^21^, with Altheim in the fifth century, showing an identical MFM of 23%, aligning well with this trend (SI Chapter S2). However, the precise geographic origins of these individuals remain unknown, as regions with similar geological conditions can be found across a broad range of locations, while sharp variations may occur within short distances ^22^, as seen in our study region between cis- and trans-Danubian territories ^21^ (SI Chapter S2). Therefore, the high MFM value could reflect short distance movements across the Danube but does not allow us to distinguish it from migration from more distant regions.

The new genetic and isotope analyses support the notion that the Limes served as a hub for people from beyond the region for centuries, facilitating exchanges across the border until the end of the Roman era and apparently continuing into the Early Medieval period ^23^. A substantial number of individuals from the North and East, described by Roman authors as “barbarians”, were integrated into the Roman army to strengthen border defences, particularly during the 4th century. The Roman fortifications along the Danube and the Rhine are thought to have been increasingly characterized by the presence of these “barbarians” ^24^. Another possible explanation for the population influx before the 5th century is the forced or voluntary migration of agricultural laborers, a phenomenon well-documented in the later Roman period. These migrants were often settled in small groups under legal restrictions as dependents of Roman landowners, which may explain the genetic homogeneity at Altheim before the collapse of Roman rule in the region during the late 5th century ^25^.

### 6th Century Spread of Romanized Populations

To investigate the origins of the IA-SE-like ancestry that emerged in southern Germany during the 6th century, we analysed genomes from the Late Roman period in Straubing-Azlburg. The site, located near several of our Early Medieval sites (Fig. 1), provides a key reference for genetic variation present in the region before the Early Medieval period. As shown in Fig. 2D, the full spectrum of the genetic diversity observed in the 6th century at the Early Medieval sites (e.g. in Altheim) was already present in the region within a Roman context before the end of Roman rule. Thus, the genetic shift we observed in the 6th century can largely be attributed to regional mobility within the broader area, including from nearby Roman settlements and former military camps. As Roman rule weakened in the late 5th century, the dissolution of military and economic structures, along with the erosion of social and legal ties between landlords and their dependent peasants — such as *coloni* and slaves — would have facilitated such local movements. While long-distance migration cannot be entirely ruled out in individual cases, it is not a necessary explanation for these patterns ^26^. Consistently, 95% of the 6th- and 7th-century individuals from Altheim can be successfully modeled in qpAdm using only 5th-century Altheim and Straubing-Azlburg sources (External_Data_Table_2.7). This is further supported by genomic analyses from other Roman camps along the Danube, such as Klosterneuburg and Ovilava, which reveal genetic diversity similar to that of 6th-century Altheim — including a distant familial link (fifth to sixth degree) with a young woman buried in Altheim (SI Chapter S6, Fig. S6.10, Extended_Data_Table3). Additionally, the distribution of genomic variation shows similarities with other sites, such as 4th century Viminacium or 6th century Szólád in Pannonia (Fig. S6.9). While similar genetic patterns in Pannonia and southern Germany could arise from mass migration between the two regions, it is more likely that they arose from parallel demographic processes, although some sporadic exchange cannot be ruled out ^8^. Indeed, the inferred contrasting social structures in these regions during the 6th century argue against significant migration between Pannonia and the Danube-Isar area ^9,13^. Furthermore, no correlation was found between the appearance of new ancestry at 6th century Altheim and non-local strontium signatures, suggesting that if long-distance migration had occurred, it came from geologically similar regions. If anything, strontium isotopes indicate a significant decrease in the proportion of non-locals in the 6th century (6%) compared to the relatively high (25%) MFM values in the 5th century in Altheim (SI Chapter S2, Extended_Data_Table1), mirroring a general trend in the wider region ^21^.

In summary, changes in the patterns of genetic variation observed from the 5th to the 6th century sites like Altheim and Büttelborn likely reflect a diffusion of ancestry from Roman foci like garrison towns and forts into the hinterland as the Empire’s control dissolved, rather than large-scale migration. Continuity in burial practices and the lack of archaeological differences between 5th and 6th century Altheim support the idea of an influx from a familiar cultural background.

### Demographic shift prompted by intermarriage between newcomers and locals

To characterize the societal impact of the post-Roman demographic shift, we reconstructed pedigrees for all individuals and traced their ancestry using a novel method that utilizes the dependencies resulting from familial relatedness to extrapolate D-statistics for unsampled individuals in a pedigree (SI Chapter S11 and S12, External_Data_Table_2.8). In Altheim, we inferred immediate intermarriage pointing to quick assimilation of incoming individuals with increased IA-SE ancestry. The rapid absorption of newcomers with a Roman background into local families of primarily northern European descent is well illustrated by the family with predominantly IA-SE ancestry located at the centre of the pedigree in Fig. 4. The four brothers of that family all had wives with D-values indicative of more IA-NC-like ancestry. This is a general trend, with females in Altheim having lower D-values than males (Wilcoxon p=0.035, Fig. S13.1). This likely reflects a continuous flow of IA-NC-like females from the northern hinterland, thereby explaining the later reduction of IA-SE-like ancestry in southern Germany, particularly if the IA-SE-like ancestry entering Altheim was primarily driven by a pulse of males from the Roman centres in the Danube-Isar region.

**Fig. 4:**
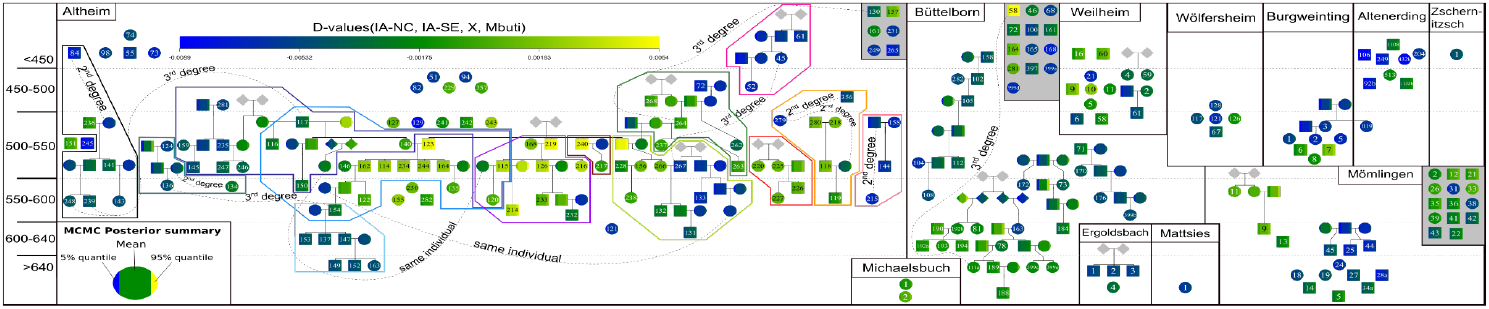
Reconstructed Pedigrees of Early Medieval sites with colour coded D-statistic values presented in a chronological context. Numbered individuals have known genomes, while unnumbered ones are inferred to have existed, with inferred D-statistic values assigned to them. Individuals not connected to a pedigree are displayed for contextual reference. The colour of the polygons surrounding Altheim family clusters matches the colour of the graves in Figure 5.

In line with the notion of a rapid assimilation of 6th-century people from a Roman background, individuals of different ancestry do not differ significantly in their grave furnishings at Altheim ^27^: We examined the grave goods of the 24 individuals with the most extreme D-values; that is, those most resembling either IA-NC or IA-SE. While clear individual variations and chronological differences in grave goods are evident, no systematic qualitative distinctions were found between IA-NC and IA-SE. However, we are currently unable to distinguish between rapid adoption of local customs by newcomers, or a pre-existing shared socio-cultural background.

### The origin of the modern European kinship system

Since Goody’s seminal study ^28^, historical and anthropological research has examined the change in family structures between Late Antiquity and the Early Middle Ages and the influence of Christianity in the emergence of the European kinship system. However, written sources for this question are limited to normative texts and anecdotal evidence on the practices of elite families. Beyond secular and ecclesiastical law and a few genomic studies on Eastern Europe ^29–32^, little is known about marriage and family practices of the rural Early Medieval population.

We used the spatial proximity of the graves in the Altheim cemetery to explore shifts in family structure. Family clusters identified in the pedigree in Figure 4 also cluster spatially in the cemetery (Fig. 5, Fig. S14.6). Statistically, first to third degree relatives were buried significantly closer together than unrelated individuals (p<0.0016), while parent-child pairs as well as siblings were buried closer together than third-degree relatives (p<0.0052, p<0.0450 respectively) (Fig. S14.1-S14.5). These findings align with IBD sharing patterns (>8 cM), which negatively correlate with burial distance (r: -0.14, p < 0.000015) and are strongest in individuals with IA-NC-like ancestry (SI Chapter S14). Furthermore, spouses — identified through shared children — were buried closer together than unrelated pairs (p < 0.0089). In Büttelborn, spouses were interred near each other and their children, with more distant relatives buried farther away (Fig. 6). While these patterns indicate a shift to a smaller family structure, our Altheim data also suggests that more distant family relationships (e.g., half-siblings) could influence burial location (see SI Chapter S14 for a detailed description of kinship patterns and the spatial distribution of relatives in the cemetery).

**Fig. 5:**
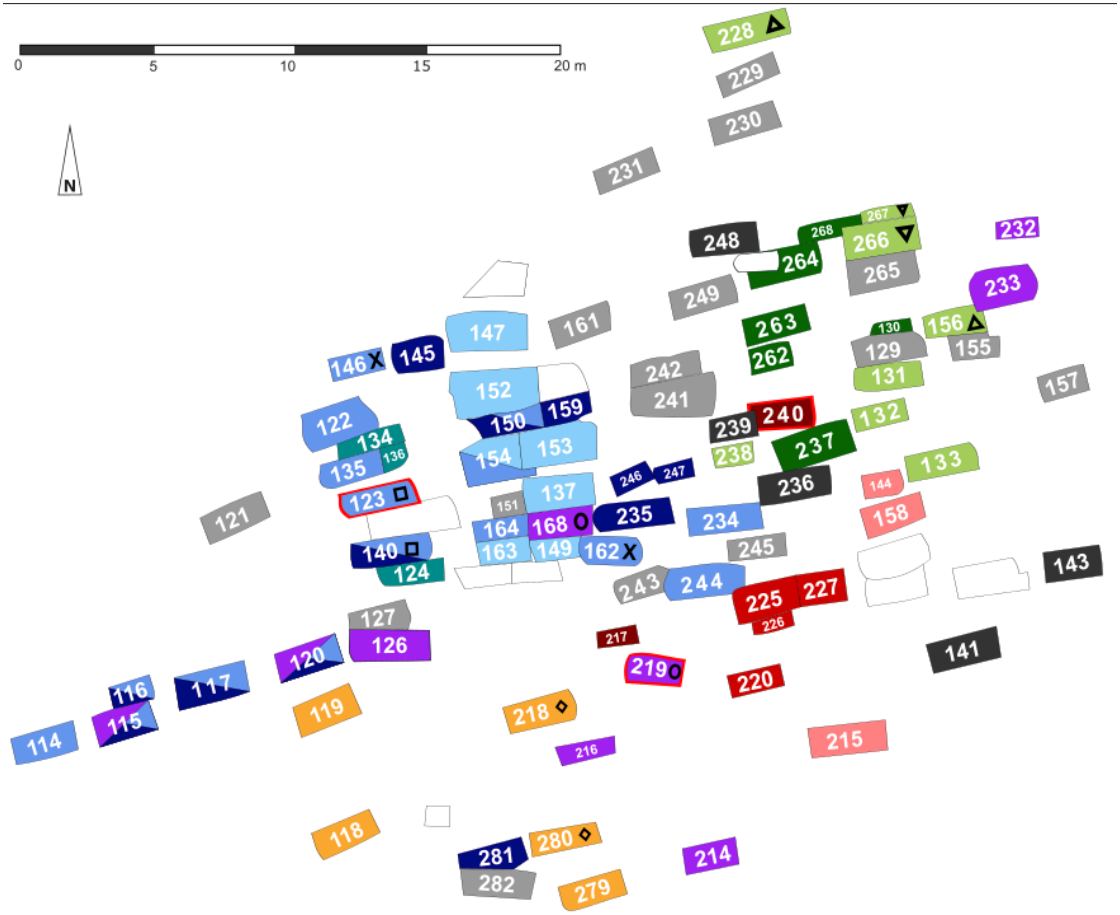
Family Clusters in the Altheim Graveyard. Position of family clusters in the north-eastern part of the Altheim graveyard (see Fig. S14.6 for the complete map). Spouses are indicated by matching symbols next to their number. Graves of unrelated individuals are grey, while unsampled ones are white. The graves of the three brothers of the central family of Fig. 4 are marked by a red border.

**Fig. 6:**
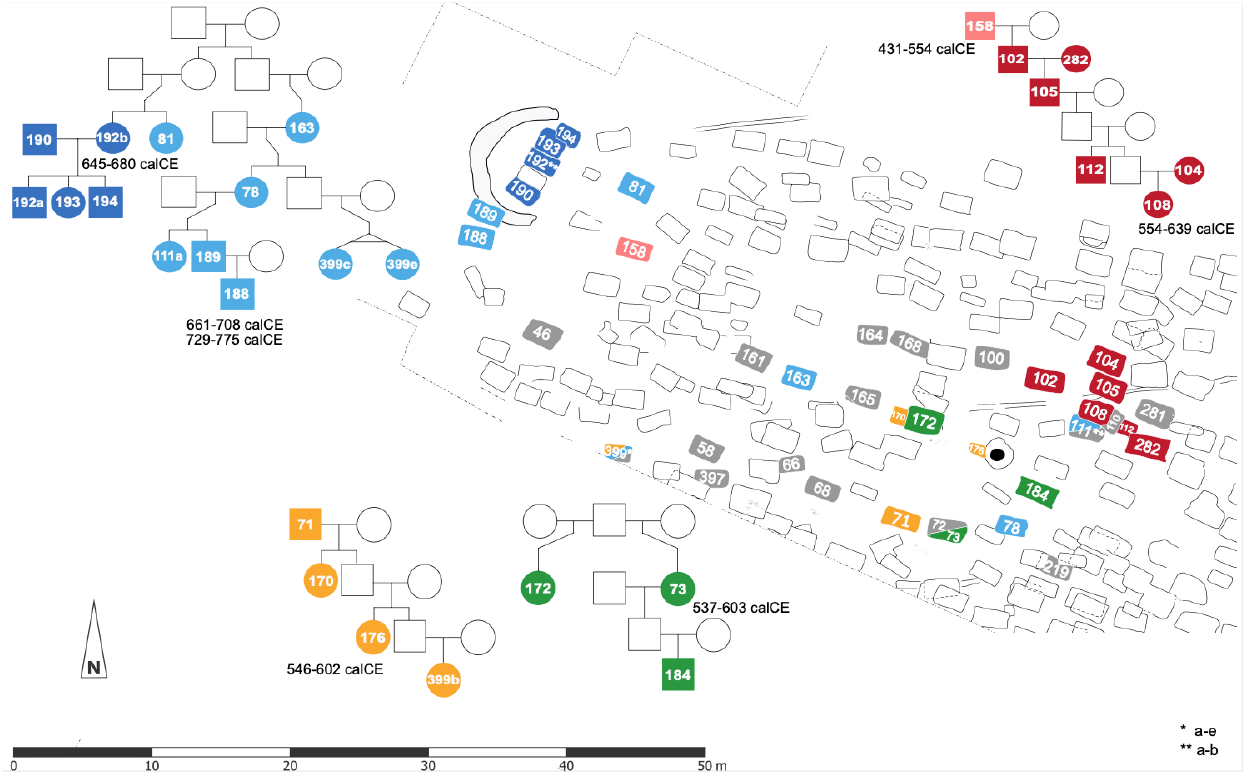
Family Clusters in the Büttelborn Graveyard. Graves of unrelated individuals are grey and unsampled ones are white. A nuclear family (father, mother, and three children, in dark blue) is buried inside the western circular structure. The red family demonstrates five generations of continuity, except for the first-generation male. While the red family (representing a male family line) shows clear clustering, the orange family (representing a female family line) shows minimal clustering.

In summary, the burial patterns indicate that the societies of Büttelborn and Altheim were based on close nuclear or stem families — sometimes complemented by kin ties such as half-siblings — reflecting a global trend of small core units within broader kin networks in simple agrarian societies ^33^ and aligning with historical research on local societies since the early Empire ^34^ and on southern Germany from c. 750 CE onwards ^35^. We do not identify any grave clusters that do not involve strong biological kinship.

Historical evidence from the Late Roman Empire suggests a shift from strict patrilineality to more flexible, bilateral inheritance systems, where male descendants typically received the larger share, but daughters and other female heirs could also inherit, and in the absence of sons, even become primary beneficiaries. By the 4th century, Roman law increasingly embraced bilateral inheritance; for example, laws permitted grandparents to pass property to the children of a deceased daughter ^34^. By the 5th and 6th centuries, both Western Roman and Justinianic law — as well as practices in western “barbarian” kingdoms — mandated equal shares for sons and daughters ^36^. Simultaneously, in Late Roman aristocratic circles, family tradition could be conceived of as deriving from both the father’s and the mother’s lines ^34^.

Drawing on our reconstructed pedigrees from the Altheim and Büttelborn graveyards, we can now investigate rules of residence and descent in non-elite agrarian societies of Central Europe. The genealogical data suggest that family lines were more frequently continued through sons (18 and 9 cases, respectively), though descent through daughters also occurred (8 and 3 cases). Notably, in three cases where descent continued through a son, an adult sister with no identified husband or children was present, while no instances of an unmarried adult brother were found when descent continued through a daughter. Women at both sites share significantly fewer IBD segments (>8 cM) than men (Altheim: 40.03 ± 7.22 vs. 170.50 ± 13.19; Büttelborn: 125.60 ± 27.05 vs. 177.74 ± 53.35), with mean relatedness coefficients 4.8 times lower in Altheim and 1.5 times lower in Büttelborn. While long-term strict patrilocality is ruled out by the high diversity in mtDNA and Y-chromosome haplogroups (Table 1, ^37^), these patterns are consistent with a loose patrilocal residence in which most women resided near their husband’s family, while some men lived near their wife’s, particularly when a son was absent. In terms of descent, both a flexible patrilineal system and a bilateral system with a preference for inheritance through sons align with these patterns. Overall, these findings suggest that residence and descent in Early Medieval southern Germany largely followed those of the Late Roman period.

In terms of marriage norms, Christian societies in the Late and post-Roman West placed increasing importance on lifelong monogamy: while divorce came under pressure from Christian rigorists, leading to stricter legal regulations, widowhood was increasingly seen as a highly esteemed status and remarriage as morally problematic ^38^. Moreover, marriages between close kin were viewed as inappropriate in Christian discourse, leading to numerous legal prohibitions issued by church councils as well as secular authorities from around 500 CE ^39^. The *Lex Baiuwariorum* (tit. VII, 1), issued in the 8th century, but reflecting earlier social norms, confirms this general pattern for Bavaria. In Altheim and Büttelborn, five individuals (four men and one woman) had multiple partners, though it remains unclear whether this reflects polygamy or serial monogamy. Nonetheless, the predominance of 61 single-partner unions suggests that monogamy was the norm in Early Medieval southern Germany. The near absence of long (>12 cM) runs of homozygosity (ROH) and the lack of shared IBD segments (>8 cM) between spouses support strict incest avoidance, excluding relationships closer than the sixth degree (Table S10.1, External_Data_Table_2.9). Given the small estimated community size (maximum of 64 individuals per settlement per generation, Table 1, SI chapter S16), incest avoidance likely promoted intermarriages across broader social networks between individuals of different ancestries. These findings indicate that normative elite texts had a tangible impact on marriage practices in lower social strata.

Recent genetic research suggests that incest avoidance was present in non-Christian societies, such as Iron Age populations from the British Isles ^40^, and Avar societies in the Western Pannonian plains ^29,41^. However, the latter shows evidence forlevirate unions, which are absent in Altheim and Büttelborn, in accordance with the *Lex Baiuwariorum*. Together, our genetic data and historical texts indicate that the family size, descent, residence, and marriage patterns prevalent in European societies until the 20th century were already present in Central Europe by the 6th century CE, most likely reflecting an adaptation of Roman elite traditions by small-scale agrarian societies, possibly as part of a broader cultural trend ^28^.

## Conclusions

While the transition from the Late Roman period to the Early Middle Ages has traditionally been framed as a conflict between northern “barbarians” and a Roman Empire in decline, recent historical, archaeological, and genetic data suggest a more nuanced process ^1,13,42–47^. In Altheim, migrants from Northern Europe, the majority of which probably had been under the influence of Roman culture for generations, established a cemetery in former Roman territories by the mid-5th century CE. At first glance, this appears to support traditional narratives. However, the layout of these cemeteries reflects rather Roman ceremonial traditions ^48^. These genetically northern migrants, building on the cultural foundation of the Roman world, developed a new burial tradition – furnishing graves with clothing, jewellery, and weapons ^1^. Rather than a simple dichotomy between “Roman” and “Germanic”, the evidence highlights a combined process of northern migration, cultural exchange, adaptation of Roman culture and the development of new social practices on the ground. To what extent this process is specific to the study region, or whether similar processes took place in other parts of the Limes region, will require further studies, particularly with genomes from Late Roman times and the early phase of the row grave horizon.

What occurs after the founding phase, however, is a distinctly transregional phenomenon in Western Europe. As Roman authority waned in the late 5th century, heightened mobility introduced a diverse population, likely originating from nearby Roman towns and military camps. In Fig. 2D, we see that the foundations of Early Medieval genetic diversity in southern Germany were already laid between the late 3rd and early 5th centuries, i.e., before the row grave horizon. During this Late Roman period, the region was home to both individuals from a non-Roman context displaying northern ancestry (sites Pförring and Kemathen) and a genetically diverse Roman military population (site Straubing-Azlburg). In Altheim, evidence of gene flow between these groups is apparent in 6th century marriages between newcomers from Roman settlements and established local families, marking a phase of integration.

Historical records align with the inferred 6th-century population shift in Altheim, with regional mobility in times of uncertainty as the favoured explanation: in the region between the Danube and Isar rivers, records indicate that Roman control began to weaken from the late 5th century onwards. Just like in Altheim and other sites in the Danube-Isar region, Büttelborn and other sites in the Rhine-Main area experienced similar gene flow from Roman settlements inside the Empire, likely driven by captivity, slavery, and military activity. The delay in demographic shifts in this region, which had already been abandoned by Roman military forces in the 3rd century, may be linked to a lesser attractiveness of an area which had not been part of the Roman Empire for more than two centuries. Further genetic analysis of human remains from nearby Roman cities like Mainz could help to clarify this. Despite these dynamic demographic shifts, core cultural practices such as monogamy, incest avoidance, and flexible inheritance systems, reflecting Late Roman norms, persisted into the Early Medieval period.

The demographic processes outlined above may serve not only as a foundational model for the persistence of cultural norms from Late Antiquity to the present but also offer a detailed framework for understanding the southward expansion of the German language in Central Europe ^19,20^. Furthermore, the interactions between northern and Late Roman populations provide valuable clues as to why Latin personal and place names, along with an early form of German, remained in use throughout the Early Middle Ages in southern Germany ^49^.

Demographic processes after the row-grave horizon remain largely unexplored. To the east of our main study region, genomic data we generated from the Austrian Molzbichl site points to an additional influx of Northern European or Baltic ancestry, potentially informing debates about the spread of Slavic languages in the medieval period ^50^. In the Danube-Isar region, the genetic composition of 7th-century populations already closely resembles those of the present day, suggesting an absence of major demographic shifts in the intervening period.

## Material and Methods

Ancient DNA extraction, library preparation, sequencing, raw-read processing and variant calling were carried out as described in Zedda *et al*. (2023) ^51^ (see SI Chapters S3 and S4 for details). Phasing and imputation of feasible genomes was performed with glimpse2 ^52,53^ for a set of sites identified as biallelic in the 1,000 genomes data set ^54^.

The birth date for each individual was estimated by subtracting the mean anthropological age-at-death estimate from the mean archaeological date estimate. Discrepancies were corrected using pedigree information, assuming a generation time of 25 years. For Büttelborn, calibrated ^14^C ranges (mean values with >93.2% probability) were used instead of archaeological dating.

SmartPCA from the EIGENSOFT package ^55^ was used for projecting ancient individuals onto a set of modern West-Eurasian individuals from the human origins data set ^56^, taken from version 54 of the Allen Ancient DNA Resource (AARD; ^57,58^). Ancient individuals were either taken directly from the AARD if available, otherwise alignments or raw-data were obtained from their respective ENA repository and processed as described in Zedda *et al*. (2023) ^51^ (see SI Chapter S4 for details).

ADMIXTOOLS version 1520 ^59^ was used for f/D-statistics and qpAdm with the following “right groups”: Mbuti.DG, Russia_Ust_Ishim.DG, CHG, EHG, Iran_GanjDareh_N, Israel_Natufian, Jordan_PPNB, Laos_Hoabinhian.SG, Russia_Samara_EBA_Yamnaya, ONG.SG, Spain_ElMiron, Turkey_N, Russia_MA1_HG.SG, Morocco_Iberomaurusian, Sunghir. For three individuals (Alh51, Alh_131, Alh_145) we had to use a different set of “right groups” (specified in External_Data_Table_2.3). For Alh_245, the individual carrying East Asian ancestry, we first carried out an f3-outgroup analysis to identify the best East Asian source populations (External_Data_Table_2.4), then added either of the three best ranking populations as potential sources (External_Data_Table_2.5).

Mitochondrial and Y-chromosomal haplotypes were determined with haplogrep3 ^60^ and Y-leaf 2 ^61^ respectively.

Runs of homozygosity were estimated with hapROH ^62^ for all genomes with at least 400K SNPs overlapping the 1240K capture array sites ^63^.

We used ancIBD ^64^ to detect identity by descent segments among our newly sequenced individuals and a curated set of previously published individuals from Europe within a 500 year range of the Early Medieval genomes, focussing on the 1240K capture array sites, as the majority of the published genomes were enriched for those sites (SI Chapter S9).

Biological relatedness was estimated using KIN ^65^ (External_Data_Table_2.10) and READ2 ^66^ (External_Data_Table_2.11) and confirmed by the IBD results (External_Data_Table_4), if possible. Pedigrees were constructed as described in Blöcher *et al*. (2023) ^67^. Correlations between the degree of relatedness and placement on the burial grounds in Altheim and Büttelborn were investigated by first overlaying the graveyard plan with a grid of arbitrary units. For each individual a point was set in the middle of their position and the coordinates were recorded so euclidean distances could be calculated. Pairs of individuals were grouped according to their degree of relatedness estimated by KIN allowing for random comparison of distances between samples from each group.

PRODAD (https://bitbucket.org/wegmannlab/prodad) was used for the pedigree-aware interpolation of individual D-statistics (see SI Chapter S12).

Individual estimates of genome-wide heterozygosity (theta) were obtained with ATLAS ^68^. Population level estimates of diversity were estimated on a set of bi-allelic transversions to minimize the effect of post-mortem damage, which is more pronounced in transitions. Hudson’s *F*_*st*_ was calculated using scikit-allele v1.3.1 (https://github.com/cggh/scikit-allel), inbreeding coefficient *F* was calculated as 1-(*Ho*/*He*), with observed (*Ho*) and expected heterozygosity (*He*) measured using plink2 ^69^. Internal (*pi*) and cross-population sequence divergence (*Dxy*) were assessed with PiXY ^70^.

SLiM ^71^ simulations were used to estimate community sizes for Büttelborn and Altheim by simulating populations with monogamous mating pairs of size N [250 - 500] distributed across n villages [2, 5], connected by migration (m - [0 - 1]) for 10 generations in 1000 replicates. In each replicate S individuals were sampled (Altheim: S=100, Büttelborn: S=40) and relatedness (r) over 3 generations was calculated. A regression analysis with the Python package (statsmodels v.0.14.4, ^72^) was used to determine the relationships between N, m and r for both communities, given the observed values (see SI Chapter S16).

## Acknowledgements

Genome data from the DZHKomics Resource with a total of 1,149 individuals, sourced from various clinics across Germany, was provided by the DZHK (German Centre for Cardiovascular Research; FKZ: 81×1500104, 81×1500302, 81×1100101) through the DZHK Heart Bank. The authors thank Sabine Schade-Lindig (hessenArchäologie), Johann Friedrich Tolksdorf (Bayerisches Landesamt für Denkmalpflege), Gerd Riedel (Stadtmuseum Ingolstadt), and Sandra Bock (Thüringer Landesamt für Denkmalpflege und Archäologie), for support. I.M. extends his gratitude to the excavation team of Viminacium for providing information regarding the burials and their dating. J. Bu. thanks Frank Siegmund for previous discussions. B.P. thanks Mihaela Jacob and Wolf-Rüdiger Teegen. Funding for genome and isotope research was provided by the Tübingen DFG Center for Advanced Studies 2496 “Migration and Mobility in Late Antiquity and the Early Middle Ages”, awarded to M.Me., S.P. and S.S.-H. Parts of this research, including the positions of L.Va., L.W. and R.M., were funded by DFG grants awarded to J.Bu. (BU 1403/19-1) and D.Q. (QU 263/3-1) and by an SNF grant to D.W. (310030_200420). M.G.T. is supported by ERC Horizon 2020 research and innovation programme grant agreements: no. 951385 (COREX) awarded to M.G.T., no. 865515 (SUSTAIN) awarded to Maria Ivanova-Bieg, no. 324202 (NeoMilk) awarded to Richard Evershed, no. 788616 (YMPACT) awarded to Volker Heyd, and by Wellcome Senior Research Fellowship Grant 100719/Z/12/Z awarded to M.G.T. M.B.’s research was funded by the German Research Foundation for his project “Co-Move” (466680522). Genetic data analyses were partially conducted using the supercomputer MOGON2 at Johannes Gutenberg University Mainz (hpc.uni-mainz.de).

## References

1. Halsall, G. Barbarian Migrations and the Roman West, 376–568. (Cambridge University Press, 2007).

2. Brownlee, E. Grave Goods in Early Medieval Europe: regional variability and decline. Internet Archaeol. (2021) doi:10.11141/ia.56.11.

3. Hamerow, H. Early Medieval Settlements: The Archaeology of Rural Communities in Northwest Europe, 400–900. (Oxford University Press, 2004).

4. Theuws, F. Long-distance trade and the Rural population of Northern Gaul. In The Oxford Handbook of the Merovingian World (eds. Effros, B. & Moreira, I.) 883–915 (Oxford University Press, Oxford, 2020).

5. Brather-Walter, S. Social ‘Mobility’ or ‘Distancing’? Spatial Organisation of Early Medieval Graveyards in South-Western Germany. In The European Countryside during the Migration Period: Patterns of Change from Iberia to the Caucasus (300-700 CE) (eds. Bavuso, I. & Castrorao Barba, A.) 201–231 (De Gruyter, 2023).

6. Brather, S. Pagan or Christian? Early medieval grave furnishings in Central Europe. In Rome, Constantinople and newly-converted Europe. Archaeological and historical evidence. Volume 1 (eds. Salamon, M.et al.) 333–349 (Kraków, Leipzig, Rzeszów, Warszawa, 2012).

7. Heitmeier, I. & Haberstroh, J. Gründerzeit: Siedlung in Bayern zwischen Spätantike und frühem Mittelalter. (EOS Verlag, 2019).

8. Veeramah, K. R. et al. Population genomic analysis of elongated skulls reveals extensive female-biased immigration in Early Medieval Bavaria. Proc. Natl. Acad. Sci. U.S.A. 115, 3494–3499 (2018).

9. Amorim, C. E. G. et al. Understanding 6th-century barbarian social organization and migration through paleogenomics. Nat. Commun. 9, 3547 (2018).

10. Gretzinger, J. et al. The Anglo-Saxon migration and the formation of the early English gene pool. Nature 610, 112–119 (2022).

11. Patterson, N. et al. Large-scale migration into Britain during the Middle to Late Bronze Age. Nature 601, 588–594 (2022).

12. Antonio, M. L. et al. Stable population structure in Europe since the Iron Age, despite high mobility. eLife 13, e79714 (2024).

13. Vyas, D. N. et al. Fine-scale sampling uncovers the complexity of migrations in 5th-6th century Pannonia. Curr. Biol. 33, 3951–3961.e11 (2023).

14. Olalde, I. et al. A genetic history of the Balkans from Roman frontier to Slavic migrations. Cell 186, 5472–5485.e9 (2023).

15. O’Sullivan, N. et al. Ancient genome-wide analyses infer kinship structure in an Early Medieval Alemannic graveyard. Sci Adv 4, eaao1262 (2018).

16. Hoffmann, J. et al. The DZHK research platform: maximisation of scientific value by enabling access to health data and biological samples collected in cardiovascular clinical studies. Clin. Res. Cardiol. 112, 923–941 (2023).

17. Gregoricka, L. A. Moving forward: A bioarchaeology of mobility and migration. J. Archaeol. Res. 29, 581–635 (2021).

18. Gretzinger, J. et al. Evidence for dynastic succession among early Celtic elites in Central Europe. Nat. Hum. Behav. 8, 1467–1480 (2024).

19. McColl, H. et al. Steppe Ancestry in western Eurasia and the spread of the Germanic Languages. bioRxiv 2024.03. 13.584607 (2024) doi:10.1101/2024.03.13.584607.

20. Speidel, L. et al. High-resolution genomic history of early medieval Europe. Nature 637, 118–126 (2025).

21. Velte, M. et al. Between Raetia Secunda and the dutchy of Bavaria: Exploring patterns of human movement and diet. PLoS One 18, e0283243 (2023).

22. Holt, E., Evans, J. A. & Madgwick, R. Strontium (87Sr/86Sr) mapping: A critical review of methods and approaches. Earth Sci. Rev. 216, 103593 (2021).

23. Geary, P. J. Before France and Germany: The Creation and Transformation of the Merovingian World. (Oxford University Press, New York, NY, 1988).

24. Behrwald, R. Gab es eine spätrömische Siedlungspolitik? in Gründerzeit: Siedlung in Bayern zwischen Spätantike und frühem Mittelalter (eds. Heitmeier, I. & Haberstroh, J.) vol. 03 447–468 (EOS Verlag, 2019).

25. Schmidt-Hofner, S. Barbarian Migrations and the Economic Challenges to the Roman Landholding Elites in the Fourth Century CE. J. late antiq. 10, 372–404 (2017).

26. Fehr, H. Friedhöfe der frühen Merowingerzeit in Baiern – Belege für die Einwanderung der Baiovaren und anderer germanischer Gruppen? in Die Anfänge Bayerns. Von Raetien und Noricum zur frühmittelalterlichen Baiovaria (eds. Fehr, H. & Heitmeier, I.) 311–336 (EOS Verlag, 2014).

27. Sebrich, J. Das spätantik-frühmittelalterliche Gräberfeld von Essenbach-Altheim. Materialhefte zur Bayerischen Archäologie 110. (Verlag Michael Laßleben, Kallmünz, 2019).

28. Goody, J. The Development of the Family and Marriage in Europe. (Cambridge University Press, Cambridge, England, 1983).

29. Wang, K. et al. Ancient DNA reveals reproductive barrier despite shared Avar-period culture. Nature 638, 1007–1014 (2025).

30. Saag, L. et al. North Pontic crossroads: Mobility in Ukraine from the Bronze Age to the early modern period. Sci. Adv. 11, eadr0695 (2025).

31. Gnecchi-Ruscone, G. A. et al. Ancient genomes reveal origin and rapid trans-Eurasian migration of 7th century Avar elites. Cell 185, 1402–1413.e21 (2022).

32. Maróti, Z. et al. The genetic origin of Huns, Avars, and conquering Hungarians. Curr. Biol. 32, 2858–2870.e7 (2022).

33. Akram-Lodhi, A. H. et al. (eds.) Handbook of Critical Agrarian Studies. (Edward Elgar Publishing, Cheltenham, England, 2021).

34. Evans-Grubbs, J. Marriage and Family Relationships in the Late Roman West. In A companion to Late Antiquity (ed. Rousseau, P.) 201–219 (Wiley-Blackwell, Oxford, UK, 2009).

35. Kohl, T. Lokale Gesellschaften: Formen der Gemeinschaft in Bayern vom 8. bis zum 10. Jahrhundert (Mittelalter-Forschungen 29). (Ostfildern, 2010).

36. Arjava, A. Women and Law in Late Antiquity. (Oxford University Press, Oxford, England, 1996).

37. Guyon, L., Guez, J., Toupance, B., Heyer, E. & Chaix, R. Patrilineal segmentary systems provide a peaceful explanation for the post-Neolithic Y-chromosome bottleneck. Nat. Commun. 15, 3243 (2024).

38. Jussen, B. Der Name der Witwe. Erkundungen zur Semantik der mittelalterlichen Bußkultur (Veröffentlichungen des Max-Planck-Instituts für Geschichte 158). (Göttingen, 2000).

39. Ubl, K. Inzestverbot und Gesetzgebung: Die Konstruktion eines Verbrechens (300-1100) (Millennium-Studien 20). (De Gruyter, Berlin, Germany, 2008).

40. Cassidy, L. M. et al. Continental influx and pervasive matrilocality in Iron Age Britain. Nature 637, 1136–1142 (2025).

41. Gnecchi-Ruscone, G. A. et al. Network of large pedigrees reveals social practices of Avar communities. Nature 629, 376–383 (2024).

42. Brather, S. Ethnische Interpretationen in der frühgeschichtlichen Archäologie. Geschichte, Grundlagen und Alternativen (Ergänzungsbände zum Reallexikon der germanischen Altertumskunde 42). (Walter de Gruyter, Berlin, 2004).

43. Brather, S. Anfang und Ende der Reihengräberfelder. Der Wandel von Bestattungsformen zwischen Antike und Mittelalter. In Antike im Mittelalter – Fortleben, Nachwirken, Wahrnehmung. 25 Jahre Forschungsverbund "Archäologie und Geschichte des ersten Jahrtausends in Südwestdeutschland” (eds. Brather, S., Nuber, H. U., Steuer, H. & Zotz, T.) 217–234 (Ostfildern, 2014).

44. Pohl, W. Introduction – Strategies of Identification: A Methodological Profile. In Strategies of Identification: Ethnicity and Religion in Early Medieval Europe (eds. Pohl, W. & Heydemann, G.) 1–64 (Turnhout, 2013).

45. Meier, M. Geschichte der Völkerwanderung. Europa, Asien und Afrika vom 3. bis zum 8. Jahrhundert n. Chr. (C H Beck, Munich, Germany, 2021).

46. Harland, J. M. Ethnic Identity and the Archaeology of the aduentus Saxonum: A Modern Framework and its Problems. (Amsterdam University Press, 2021).

47. Tian, Y. et al. The role of emerging elites in the formation and development of communities after the fall of the Roman Empire. Proceedings of the National Academy of Sciences 121, e2317868121 (2024).

48. Fehr, H. Die Anfänge des Reihengräberhorizontes: Archäologische Aspekte. Germanen und Romanen im Merowingerreich. Frühgeschichtliche Archäologie zwischen Wissenschaft und Zeitgeschehen. Ergänzungsbände zum Reallexikon der Germanischen Altertumskunde. vol. 68 725–783 (De Gruyter, Berlin, Germany, 2010).

49. Haubrichs, W. The Multilingualism of the Early Middle Ages: Evidence from Peripheral Regions of the Regnum orientalium Francorum. In The Languages of Early Medieval Charters. Latin, Germanic Vernaculars and the Written Word (eds. Gallagher, R., Roberts, E. & Tinti, F.) 68–116 (Leiden/Boston, 2021).

50. Stolarek, I. et al. Genetic history of East-Central Europe in the first millennium CE. Genome Biol. 24, 173 (2023).

51. Zedda, N. et al. Biological and substitute parents in Beaker period adult-child graves. Sci. Rep. 13, 18765 (2023).

52. Rubinacci, S., Hofmeister, R., da Mota, B. S. & Delaneau, O. Imputation of low-coverage sequencing data from 150,119 UK Biobank genomes. bioRxiv 2022.11.28.518213 (2022) doi:10.1101/2022.11.28.518213.

53. Rubinacci, S., Ribeiro, D. M., Hofmeister, R. J. & Delaneau, O. Efficient phasing and imputation of low-coverage sequencing data using large reference panels. Nat. Genet. 53, 120–126 (2021).

54. 1000 Genomes Project Consortium et al. A global reference for human genetic variation. Nature 526, 68–74 (2015).

55. Patterson, N., Price, A. L. & Reich, D. Population structure and eigenanalysis. PLoS Genet. 2, e190 (2006).

56. Lazaridis, I. et al. Ancient human genomes suggest three ancestral populations for present-day Europeans. Nature 513, 409–413 (2014).

57. Mallick, S. & Reich, D. The Allen Ancient DNA Resource (AADR): A curated compendium of ancient human genomes. Harvard Dataverse 10.7910/DVN/FFIDCW (2023).

58. Mallick, S. et al. The Allen Ancient DNA Resource (AADR) a curated compendium of ancient human genomes. Sci. Data 11, 182 (2024).

59. Patterson, N. et al. Ancient admixture in human history. Genetics 192, 1065–1093 (2012).

60. Schönherr, S., Weissensteiner, H., Kronenberg, F. & Forer, L. Haplogrep 3 -an interactive haplogroup classification and analysis platform. Nucleic Acids Res. 51, W263–W268 (2023).

61. Ralf, A., González, D. M., Zhong, K. & Kayser, M. Yleaf: Software for Human Y-Chromosomal Haplogroup Inference from Next-Generation Sequencing Data. Molecular Biology and Evolution 35, 1291–1294 (2018).

62. Ringbauer, H., Novembre, J. & Steinrücken, M. Parental relatedness through time revealed by runs of homozygosity in ancient DNA. Nat. Commun. 12, 5425 (2021).

63. Mathieson, I. et al. Genome-wide patterns of selection in 230 ancient Eurasians. Nature 528, 499–503 (2015).

64. Ringbauer, H. et al. Accurate detection of identity-by-descent segments in human ancient DNA. Nat. Genet. 56, 143–151 (2024).

65. Popli, D., Peyrégne, S. & Peter, B. M. KIN: a method to infer relatedness from low-coverage ancient DNA. Genome Biol. 24, 10 (2023).

66. Alaçamlı, E. et al. READv2: advanced and user-friendly detection of biological relatedness in archaeogenomics. Genome Biol. 25, 216 (2024).

67. Blöcher, J. et al. Descent, marriage, and residence practices of a 3,800-year-old pastoral community in Central Eurasia. Proc. Natl. Acad. Sci. U.S.A. 120, e2303574120 (2023).

68. Link, V. et al. ATLAS: Analysis Tools for Low-depth and Ancient Samples. bioRxiv 105346 (2017) doi:10.1101/105346.

69. Chang, C. C. et al. Second-generation PLINK: rising to the challenge of larger and richer datasets. GigaScience 4, 7 (2015).

70. Korunes, K. L. & Samuk, K. pixy: Unbiased estimation of nucleotide diversity and divergence in the presence of missing data. Mol. Ecol. Resour. 21, 1359–1368 (2021).

71. Haller, B. C. & Messer, P. W. SLiM 3: Forward genetic simulations beyond the Wright-Fisher model. Mol. Biol. Evol. 36, 632–637 (2019).

72. Seabold, S. & Perktold, J. Statsmodels: Econometric and Statistical Modeling with Python. In Proceedings of the 9th Python in Science Conference (eds. van der Walt, S. & Millman, J.) 92–96 (2010).

